# Prebiotic ALPINA GOS produced from whey permeate has a bifidogenic effect on the adult fecal microbiota *in vitro*, including stimulation of organic acids production

**DOI:** 10.64898/2026.01.26.701770

**Authors:** David Orrego, Guus A.M. Kortman, Eric R. Hester, Laura Sierra-Zapata, Shadia Blel-Jubiz, Vanesa Caro-Miranda, Bernadette Klotz-Ceberio

## Abstract

Functional oligosaccharides, such as galacto-oligosaccharides (GOS), are valued for modulating gut microbiota and promoting health. This study aimed to produce a high-purity GOS ingredient (ALPINA GOS) via nanofiltration/diafiltration and assess its prebiotic efficacy using an in vitro fermentation model. GOS-rich syrup was obtained from transgalactosylation of lactose in concentrated whey permeate (30% lactose) and processed by diafiltration/nanofiltration to reduce monosaccharides and enrich oligosaccharide content. Carbohydrate composition was analyzed by HPAEC-PAD. Prebiotic activity was evaluated using a MicroColon model with fecal inocula from healthy adults, measuring pH, short-chain fatty acids (SCFAs), and microbiota shifts. Membrane processing increased oligosaccharides from 55.5% to 70.2% (dry basis) and reduced monosaccharides from 25.2% to 5.1%. ALPINA GOS induced a dose-dependent pH reduction and significantly enhanced lactate and acetate production, with stronger effects at 10 mg/mL. Microbiota profiling showed increased abundance of beneficial bacteria, especially *Bifidobacterium*, versus control. The findings confirm that GOS can be sustainably produced from whey permeate and exhibits potent prebiotic activity, supporting its application in functional foods aimed at gut health.

## 1. Introduction

Functional oligosaccharides are non-digestible carbohydrates that can exhibit prebiotic activity when metabolized by the human gut microbiota. Notably, they selectively promote the growth of beneficial bacteria, such as bifidobacteria and lactobacilli, which can trigger host health benefits (Mei et al., 2022). Galacto-oligosaccharides (GOS) are well-known prebiotics derived primarily from lactose. GOS consists of a glucose moiety followed by several galactose molecules forming an oligomer with 2 to 10 units. These units are connected by β (1-2, 1-3, 1-4, 1-6) or α (1-6) linkages, depending on the source or production process (Wang et al., 2023). Specifically, β-GOS are produced through the transgalactosylation activity of β-galactosidase which is favored by high lactose concentration and limited water availability (Mei et al., 2022). Under these conditions, the enzyme preferentially catalyzes the transfer of galactose to hydroxyl groups on other lactose molecules, rather than hydrolyzing lactose into glucose and galactose, resulting in GOS formation (Tzortzis & Vulevic, 2009).

Commercial GOS ingredients are typically produced from milk-derived lactose by enzyme treatment. However, the increasing interest in circular economy principles has driven the utilization of industrial by-products, such as cheese whey, which contains significant amounts of lactose derived from milk. To optimize enzyme activity and maximize GOS production, a reduction in water content is recommended, achieving a lactose concentration of at least 30% w/w. Fischer and Kleinschmidt (Fischer & Kleinschmidt, 2018) observed that GOS yields from the enzymatic conversion of lactose using β-galactosidase from *Aspergillus oryzae* reach a peak at 30% lactose, maintaining a plateau up to 60% lactose concentration.

Cheese whey contains most of the lactose in milk, as well as whey protein, fat, and some of the minerals and vitamins in milk (Roy et al., 2020). This whey can be processed to separate its components; for example, fat is removed using a cream separator, and whey protein is extracted via ultrafiltration (UF), leaving a retentate rich in protein, and a permeate containing lactose and minerals (Bylund, 1995). The lactose in the permeate can be further concentrated through nanofiltration (NF), which removes water and monovalent minerals, resulting in a retentate suitable for GOS production (Chen et al., 2023). The use of whey permeate as a substrate for GOS synthesis not only enhances the functional properties of whey but also generates prebiotic compounds known for their beneficial effects on gut health, (Fischer & Kleinschmidt, 2018; Souza et al., 2022). Thus, leveraging cheese whey for GOS production addresses environmental concerns and capitalizes on the growing demand for functional foods and nutraceuticals.

The enzymatic reaction used to produce GOS from lactose results in the formation of oligosaccharides, but carbohydrates like lactose, glucose, and galactose remain in the reaction mix. To purify and concentrate GOS, various methods are employed, including chromatography, vacuum distillation, and nanofiltration (Olivares-Tenorio et al., 2022). Traditional methods like cation exchange chromatography and vacuum distillation require substantial time, water, and energy, making them less practical for industrial-scale applications (Nobre et al., 2009). In contrast, nanofiltration offers a more efficient alternative, capable of handling high solute concentrations with lower energy requirements. Research has shown that nanofiltration membranes, particularly those made from polyethersulfone, maintain high flux and selectively reject undesired sugars under varying pressures, achieving high-purity GOS (Schmidt et al., 2019). These strategies to concentrate GOS by removing digestible sugars allow to obtain an ingredient with lower glycemic index and enhanced functionality that can be translated to the final consumer

*In vitro* and *in vivo* studies consistently demonstrate that prebiotic carbohydrates, whether consumed in food or supplements, are fermented by members of the microbiota in the large intestine, producing short-chain fatty acids (SCFAs), methane, hydrogen, carbon dioxide, and other metabolites (Liu et al., 2022; Slavin, 2013). These metabolites play a crucial role in promoting intestinal health and human health in general via metabolic, immunological and neurological pathways (Krishnamurthy et al., 2023; Roager & Licht, 2018). SCFAs are the primary byproducts of fiber fermentation by gut bacteria and contribute to the integrity of the intestinal barrier by fueling colonocytes and enhancing tight junctions between cells (Portincasa et al., 2022). Consumption of SCFA) by colonocytes contributes to a hypoxic environment in the gut lumen, as these cells require increased oxygen to oxidize SCFAs for energy production. For instance, butyrate derived from bacteria enhances epithelial oxygen consumption, which stabilizes hypoxia-inducible factor (HIF), a transcription factor that coordinates barrier protection (Kelly et al., 2015; Litvak et al., 2018).

Furthermore, SCFAs exhibit anti-inflammatory properties and promote the regulation of immune responses, which helps to maintain a balanced microbiota and prevent gut dysbiosis (Venegas et al., 2019; Zhang et al., 2022). Additionally, SCFAs have been shown to alleviate allergic skin reactions, to balance the central nervous system, regulate genetic and epigenetic mechanisms, and to improve calcium solubility and absorption due to lowered colonic pH (Canani et al., 2011; Griffin et al., 2002; Hong et al., 2015; Mei et al., 2022; Senchukova, 2023). Despite these known benefits, the intricate mechanisms underlying the interactions within the intestinal ecosystem in both health and disease contexts are still being extensively researched (Canani et al., 2011). These combined effects underscore the importance of fiber in ensuring a healthy gut ecosystem and overall intestinal function (Kaur et al., 2024; Pathan et al., 2024). As a prebiotic ingredient, GOS modulates the gut microbiota by specifically promoting the growth of *Bifidobacterium* and, to a lesser extent, *Lactobacillus*. These bacteria play a crucial role in the production of lactic acid and SCFAs such as acetate in the colonic lumen (Sangwan et al., 2011). The health benefits associated with GOS are primarily attributed to its fermentation and the resulting production of SCFAs, along with the selective growth of beneficial gut bacteria (Grimaldi et al., 2016).

This study underscores the multifaceted advantages of using a GOS-rich syrup derived from whey permeate. By converting a dairy by-product into a valuable prebiotic ingredient, the research addresses both its environmental sustainability and circularity, as well as its health promotion effects. The production of GOS from whey permeate not only mitigates waste but also taps into the expanding market for functional foods that support gut health, with relevant evidence that supports these effects. The innovative application of nanofiltration technology ensures high purity and efficiency, making this approach feasible for large-scale industrial adoption. In addition, the *in vitro* prebiotic activity of the novel GOS ingredient produced through the described process was evaluated using a MicroColon fermentation model (Kortman et al., 2023). This assessment examined its effects on medium pH, short-chain fatty acid production, and the microbiota derived from human fecal donors. Overall, the study provides a comprehensive framework and methodology for testing a sustainable method of producing GOS and evaluating its efficacy.

## 2. Materials and Methods

### 2.1 Production and concentration of the GOS syrup

To produce the GOS syrup, concentrated whey permeate (CWP) was obtained from the tangential filtration plant of ALPINA Productos Alimenticios S.A.S. BIC, located in Sopó, Cundinamarca (Colombia). CWP at 20% w/w lactose was further concentrated through vacuum evaporation until 30% lactose was reached. Evaporated CWP (E-CWP) was used as a substrate for the enzymatic production of a GOS-rich syrup as described by Orrego and Klotz-Ceberio (2022) (Orrego & Klotz-Ceberio, 2022). Table 1 presents the composition of the GOS syrup.

**Table 1.**
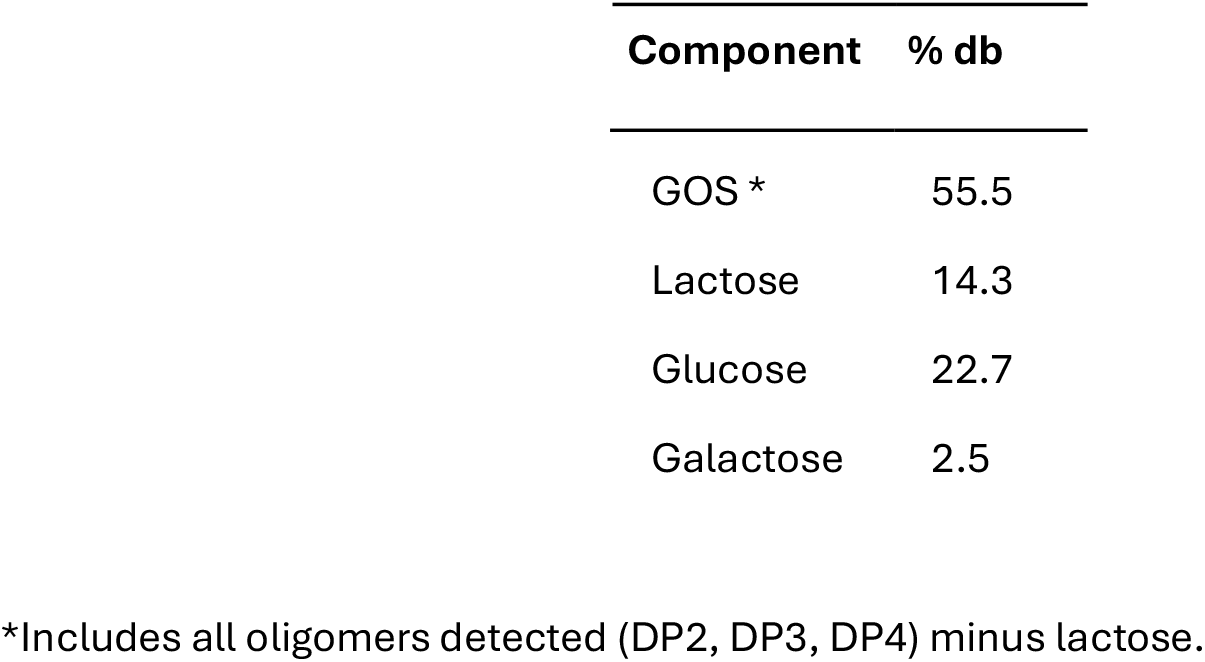
Carbohydrate composition of the GOS syrup.

To increase the GOS content and reduce the concentration of digestible sugars in the syrup, a diafiltration/nanofiltration (DF/NF) strategy was implemented. The GOS syrup was 10 times diluted before filtration. The GOS solution was concentrated batch-wise to a concentration factor (CF) of 2 at 55°C and 10 bar using a lab20 system (Alfa Laval) with NFX membranes (150-300Da; Synder filtration). Next, the concentrate was diafiltered batch-wise to 500%. For analysis of sugars and GOS, samples were taken of the syrup, diluted syrup, concentrates and permeates at CF 2, after 100, 300 and 500% diafiltration. Before application in MicroColon, the syrup was further concentrated 3.2 times on a rotary evaporator. The concentrated syrup was identified as ALPINA GOS throughout the manuscript.

### 2.2 Sugar and GOS analysis by HPAEC-PAD

High Performance Anion Exchange Chromatography with Pulsed Amperometric Detection (HPAEC-PAD) on a gold electrode was used for the quantitative analyses of GOS. The analyses were performed with a ICS-5000 DP pump, AS-AP autosampler, DC column compartment and ED electrochemical detector (Thermo Scientific).

To 0.1 g GOS solution, 9.9 g of a 10% acetonitrile solution was added and the solution was mixed well. After centrifugation 5 minutes at 14 000 g, 200 µL filtrate was transferred to an HPLC vial with insert. 5 mL sample was injected on a Carbopac PA-1, 250 × 2 mm, column (Thermo Scientific) thermostated at 30°C. The galacto-oligosaccharides were eluted at a flow rate of 0.25 mL/min with a linear gradient of 44 mM sodium hydroxide + 10 mM sodium acetate to 76 mM sodium hydroxide + 80 mM sodium acetate in 48 minutes.

Data analysis was done with Chromeleon software version 7.3.1 (Thermo Scientific). Quantitative analyses were carried out using standard solutions of the mono and oligosaccharides (Merck) and Vivinal GOS (Friesland Campina). The identification of the different GOS component was done according to Coulier et al. (2009) (Coulier et al., 2009).

### 2.3 Prebiotic activity Measurement

#### 2.3.1 Chemicals and test compounds

Chemicals were purchased from Merck Life Science Germany GmbH (Darmstadt, Germany), unless stated otherwise. Short-chain fructo-oligosaccharides (scFOS) (Orafti P95) was purchased from Beneo (Mannheim, Germany). 10x concentrated stock solutions of scFOS, ALPINA GOS syrup and the sugar control (glucose, lactose and galactose) were made by dissolving in reverse osmosis (RO) water and subsequent filter sterilization.

ALPINA GOS and scFOS were assessed at final concentrations of 5 and 10 mg/mL. Based on the sugar composition of ALPINA GOS, a sugar control was included, which was composed of 0.35 or 0.7 mg/mL glucose, 0.03 or 0.06 mg/mL galactose and 1.75 or 3.5 mg/mL lactose, respectively (mono and disaccharide mix, called “MonoDiMix”). The non-treated control was gut-like medium without addition of ingredients (water control).

#### 2.3.2 Recruitment of healthy donors and fecal sample preparation

Fecal samples were collected from 4 healthy, non-smoking, adult volunteers (Dutch nationality) with the following characteristics: age 22-66 years, 3 females, 1 male, BMI 21.2-25.2, omnivore diets. None of the volunteers ingested antibiotics for at least six months prior to feces collection, nor did they consume laxatives and did not consume pro- and prebiotics for at least two weeks prior to feces collection (yoghurt and cheeses were permitted). The volunteers were not pregnant at the time of feces collection. Each volunteer was assigned a number (27, 28, 29, or 30) to be identified in the results section. Written informed consent was obtained from all participants.

The volunteers collected their feces using a stool collection kit. The collection kit contained a Fecotainer (Excretas Medical B.V., The Netherlands), sterile sampling spoon, BD GasPak EZ Pouch System bag (New Jersey, USA), BD GasPak EZ Pouch System with indicator (New Jersey, USA), 50 mL Cellstar Cellreactor tube (Greiner bio-one, Frickenhaus, Germany), and a degradable plastic sheet to place into the Fecotainer. A subsample of fresh feces was transferred to the 50 mL Cellstar Cell reactor tube after which it was carefully sealed in the BD GasPak EZ Pouch System bag together with a BD GasPak EZ Pouch System with indicator. The fecal samples were kept anaerobic and refrigerated, and were transferred to NIZO food research B.V. (The Netherlands) within 16h to an anaerobic chamber (5% CO_2_, 5% H_2_, 90% N_2_) to prepare the fecal inoculates. The fecal inoculates were prepared by diluting the stool to 10% (w/v) using 1M anaerobic phosphate buffered saline (PBS) and were vortexed to homogenize the feces. The resulting fecal slurries from individual volunteers were used immediately as inoculum for the MicroColon experiment.

#### 2.3.3 MicroColon batch fermentation experiment

The NIZO MicroColon model is an anaerobic batch fermentation model based on a 96-well plate format. Using human fecal samples as inoculum and gut-like culture media, the model reproducibly simulates the microbial fermentation processes in the human colon. This high-throughput screening model is being used to evaluate the effect of ingredients on gut microbiota composition and metabolism.

A low-carbohydrate gut-like medium (LCM), based on the medium of Macfarlane et al. (Macfarlane et al., 1998) and Kim et al. (Kim et al., 2011) was used. The LCM contained per liter: 0.5 g starch (potato), 4 g porcine gastric type III mucin, 0.5 g pectin (citrus), 0.5 g guar gum, 0.5 g xylan (birchwood), 0.5 g arabinogalactan (larch wood), 3 g casein, 0.5 mL peptone water, 0.5 g tryptone, 0.4 g bile salt, 4.5 g yeast extract, 5 mg FeSO4•7H2O, 4.5 g NaCl, 4.5 g KCl, 0.5 g KH2PO4, 0.5 g K2HPO4, 1.25 g MgSO4•7H2O, 0.1 g CaCl2•2H2O, 1.5 g NaHCO3, 0.8 g l-cysteine HCl (anhydrous), 1 mL Tween-80, and 1 mL hemin. The hemin stock solution was prepared by adding a few drops of NaOH before dissolving in milli-q water. After autoclaving for 15 min at 121°C, the culture medium was placed anaerobically for at least 12 hours prior to inoculation. The fecal samples 10% (w/v) were inoculated in the culture medium to a final concentration of 0.1% w/v. Next, 900 µL inoculated culture medium was added to the wells of 96-deep-well plates containing 100 µL of the 10x concentrated test ingredients, or water control. The plates were sealed using a sterile Silverseal (Greiner bio-one, Frickenhaus, Germany), and were statically incubated in the anaerobic chamber at 37°C for 6 and 20 hours.

At harvesting, pH was measured for each of the wells using a micro (7 mm diameter) pH electrode (Loveland, USA) with cleaning in between. Next, the cell suspensions were transferred to 1.5 mL tubes and centrifuged at 14000 rpm for 10 min. The pellets were stored at -20°C for later DNA extraction and the supernatant was stored until analysis of SCFA content.

#### 2.3.4 SCFA analysis by HPLC

For organic acid analysis, samples were prepared according to a modified and previously described method (Gommers et al., 2019). 100 µL of MicroColon supernatant sample was diluted with 1 ml of 1M perchloric acid (HClO4) to extract the organic acids. Lipids and proteins in the fecal sample were removed by centrifugation for 5 min at 20,000 g. Organic acids lactate, acetate, propionate, butyrate, formate and succinate were determined by high-performance anion-exchange chromatography with UV and refractive index detection. 25 µl of the supernatant was injected on a guard column in series with 2 Rezex ROA-Organic Acid H+ Analytical Columns (Phenomenex, Torrance, CA, USA). The organic acids were eluted in isocratic mode with 5 mM sulfuric acid (H_2_SO_4_) with a flow rate of 0.60 ml/min. The column oven was held at a temperature of 60°C. Data analysis was performed with Chromeleon software v.7.2 (Thermo Fisher Scientific). The result was calculated using 5 concentrations of a standard mixture containing all relevant organic acids, which was used as a reference sample in each continuous series of analysis.

#### 2.3.5 DNA extraction for microbiota characterization

Cell pellets were thawed at 4°C and suspended in 700 µL S.T.A.R. buffer (Roche, Indianapolis, IN, USA). The suspension was transferred to 0.5 g of 0.1 mm sterilized zirconia beads in a 2.0 mL screw-cap tube. Lysis was performed in a FastPrep instrument (MP Biomedicals, Santa Ana, CA, USA) at 5.5 ms for 3 times 1 min at room temperature. Samples were then incubated, while shaking at 100 rpm, at 95°C for 15 min, after which samples were centrifuged at 16000 g for 5 min at 4°C. The supernatant was collected and kept on ice. The pellet was subjected to another lysis round as described above, except only 350 µL S.T.A.R. buffer was added. The resulting supernatant was pooled with the supernatant of the first lysis round. Purification of DNA was performed on the automated Maxwell instrument (Promega, Madison, WI, USA) by applying the Maxwell 16 Tissue LEV Total RNA Purification Kit (Promega) according to the manufacturer’s instructions. 250 µL of the supernatant was applied and 50 µL of RNAse/DNAse free water was used for elution of the DNA.

#### 2.3.6 Microbiota composition profiling by 16S rRNA gene sequencing

For 16S rRNA sequencing, Barcoded amplicons from the V3-V4 region of 16S rRNA genes were generated using a 2-step PCR, and universal primers appended with Illumina adaptors were used for initial amplification of the V3-V4 part of the 16S rRNA gene with the following sequences: forward primer, ‘5-CCTACGGGAGGCAGCAG-3’ (broadly conserved bacterial primer 357F); reverse primer, ‘5-TACNVGGGTATCTAAKCC’ (broadly conserved bacterial primer (with adaptations) 802R) appended with Illumina adaptor sequences. PCR amplification mixture contained: 2 µL 10x diluted sample DNA, 1 µL bar-coded forward primer, 15 µL master mix (1 µL KOD Hot Start DNA Polymerase (1 U/µL; Novagen, Madison, WI, USA), 5 µL KOD-buffer(10×), 3 µL MgSO4 (25 mM), 5 µL dNTP mix (2 mM each), 1 µL (10 µM) of reverse primer) and 32 µL sterile water (total volume 50 µL). PCR conditions were: 95°C for 2 min followed by 35 cycles of 95°C for 20 sec, 55°C for 10 sec, and 70°C for 15 sec. Size of the amplicons was checked using agarose gel electrophoresis. The approximately 500 bp PCR amplicon was subsequently purified using the MSB Spin PCRapace kit (Invitek, Berlin, Germany) and concentration was checked with a Quant-IT dsDNA Assay kit (Thermo Fisher Scientific) and microplate spectrophotometer. For the library PCR step with sample-specific barcoded primers, purified PCR products were shipped to BaseClear BV (Leiden, The Netherlands). PCR products were checked on a Bioanalyzer (Agilent) and quantified. This was followed by multiplexing, clustering and sequencing on an Illumina MiSeq with the paired-end (2x) 300 bp protocol and indexing. FASTQ read sequence files were generated using bcl2fastq version 2.20 (Illumina). Initial quality assessment was based on data passing the Illumina Chastity filtering. Subsequently, reads containing PhiX control signal were removed using an in-house filtering protocol. In addition, reads containing (partial) adapters were clipped (up to a minimum read length of 50 bp). The second quality assessment was based on the remaining reads using the FASTQC quality control tool version 0.11.8. (http://www.bioinformatics.babraham.ac.uk/projects/fastqc/).

#### 2.3.7 Microbiome analysis and statistics

Bioinformatics analyses on the sequencing data were done with a NIZO workflow utilizing amplicons from the V3-V4 16S rRNA region. The workflow incorporated FastQC and multiQC as quality filtering and assessment of raw sequencing reads. ASVs were then generated using DADA2, and the taxonomic assignment of ASVs was performed using a naive Bayesian classifier (using SILVA v138; (Callahan et al., 2016; Quast et al., 2013)). The species-level taxonomy was assigned based on exact sequence matching (100% identity). ASVs were aligned using MAFFT, and a phylogenetic tree was produced using FastTree (Katoh et al., 2009; Price et al., 2010). Counts were rarefied to 11,342 counts/sample. Alpha diversity was evaluated using the Shannon index, which accounts for both richness and evenness of bacterial communities, and Faith’s phylogenetic diversity, which considers the phylogenetic breadth of the communities. These metrics were calculated with the vegan and picante packages in R (Dixon, 2003; Kembel et al., 2010). Beta diversity was assessed using the weighted UniFrac distance metric, which incorporates phylogenetic information and accounts for the relative abundances of taxa. Principal coordinates analysis (PCoA) plots were generated with the ggplot2 package to visualize group differences. All visualizations were created using the ggplot2 package in R software (Lozupone & Knight, 2005; R Core Team, 2023; Wickham, 2011).

#### 2.3.8 Statistical analysis

Given the small sample size (n = 4), the statistical analyses are exploratory. The false discovery rate (FDR) correction for multiple testing was applied using the Benjamini-Hochberg method for all taxa, except for *Bifidobacterium* and *Lactobacillaceae*, as for these two taxa of primary interest an effect was expected. To evaluate the effect of the ingredients on the production of each organic acid, at each time point, Friedman test with Dunn’s posthoc test was applied. This was also done on the sum (total) of organic acids (lactate + acetate + butyrate + propionate + succinate + formate). To test the effect of ALPINA GOS on microbiota composition of each volunteer a PERMANOVA analysis was done. Finally, to create the Principal Component Analysis (PCoA) of the microbiome of the 4 donors, Weighted UniFrac distances in R using the vegan package were calculated; data for this analysis can be found in Supplementary Material 2 (SM2). Differences in taxa relative abundance across treatments were tested for significance using the Friedman test with a Dunn’s posthoc test using the rstatix package implementation (Kassambara, 2019).

## 3. Results and Discussion

### 3.1 Nanofiltration and diafiltration of the GOS syrup allows to increase GOS and reduce monosaccharide concentration

As a result of the transgalactosylation reaction used to produce GOS from lactose, significant amounts of digestible sugars, such as glucose and lactose, are commonly found. These sugars, particularly glucose, are considered undesirable due to their higher caloric content (4 kcal/g) and glycemic index (100) (Shkembi & Huppertz, 2023), compared to GOS, which has a caloric content below 2 kcal/g and a glycemic index of 0 (Van Leusen et al., 2014).

Furthermore, the presence of lactose in such ingredients limits their application in lactose-free products and may result in rejection by lactose-intolerant consumers.

The GOS syrup developed by Orrego & Klotz-Ceberio (2022) (Orrego & Klotz-Ceberio, 2022) represents an innovative approach to valorizing whey permeate by generating a prebiotic fiber. However, the initial formulation contains a significant amount of digestible monosaccharides and lactose (see Table 1), which could be reduced or eliminated using various methods.

Several techniques have been employed to concentrate GOS while removing glucose, galactose, and lactose, including chromatographic separation, membrane fractionation, selective fermentation, precipitation, adsorption, and electrodialysis (Olivares-Tenorio et al., 2022). Among these technologies, membrane fractionation was selected for this study due to its cost-efficiency and scalability in food companies with tangential filtration capabilities.

However, concentrating GOS based on molecular weight presents challenges, particularly in removing lactose, as GOS with a degree of polymerization (DP) of 2 has the same molecular weight as lactose. Therefore, lactose removal can significantly reduce the GOS concentration in the ingredient. To address this, our efforts in this study focused on removing monomeric sugars such as glucose and galactose.

Table 2 presents the carbohydrate composition of the GOS syrup before and after diafiltration/nanofiltration. As shown, the oligosaccharide concentration increased from 55.5% to 70.2%, while allolactose + lactose rose from 19.3% to 24.7%. Conversely, the monosaccharide content (glucose and galactose) decreased from 25.2% to 5.1%, consistent with the use of a nanofiltration membrane with a molecular weight cut-off (MWCO) of 300 Da.

**Table 2.**
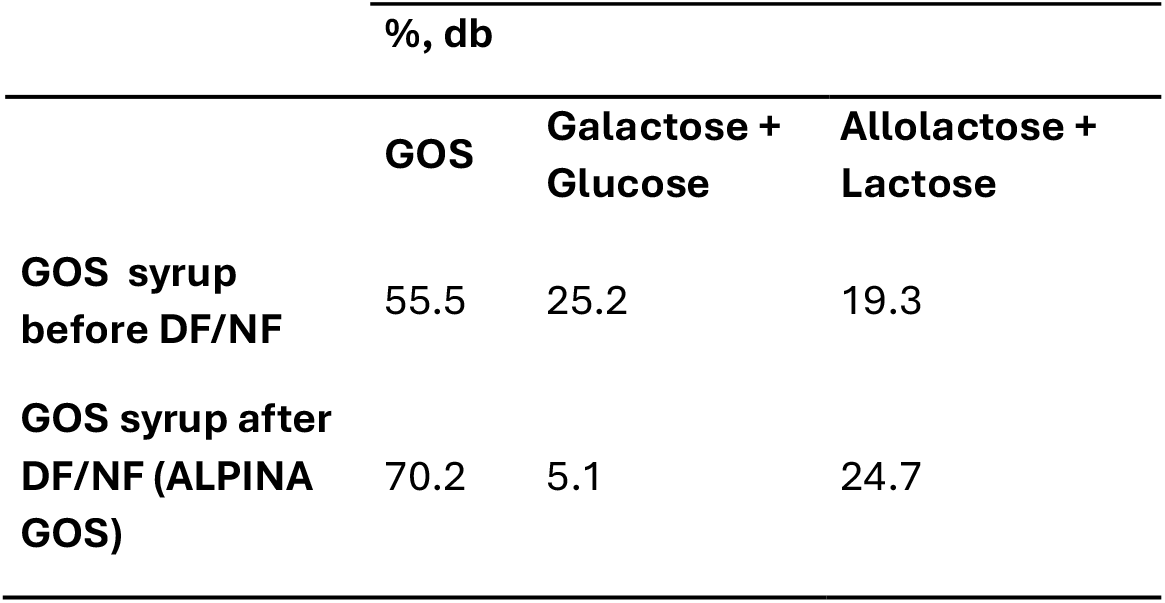
Total GOS, Mono and Disaccharides after DF/NF treatment.

Although the total elimination of digestible sugars is ideal, a 20.1% reduction in monosaccharides and a 14.7% increase in fiber significantly enhance the nutritional value of the resulting ingredient. The composition of the ingredient obtained after the DF/NF process is comparable to commercial GOS ingredients, such as Biotis GOS-P (FrieslandCampina), Oligomate 55NP (Yakult Pharmaceutical Industry Co., Ltd), and Cup Oligo P (Nissin Sugar Co., Ltd), according to the compositions reported by Olivares-Tenorio et al. (2022) (Olivares-Tenorio et al., 2022).

Several studies have evaluated GOS concentration using NF membranes with various MWCOs (Córdova et al., 2016; Feng et al., 2009; Michelon et al., 2014). For instance, Michelon et al. (2014) (Michelon et al., 2014) found that a 400 Da membrane had a rejection coefficient (RC) of 0.61 for GOS ≥DP3, 0.25 for GOS DP2 and lactose, 0.22 for glucose, and 0.23 for galactose, allowing some GOS to pass through. However, using membranes with lower MWCOs might improve rejection coefficients for GOS, while removing some monosaccharides. In our study, applying a 150–300 Da NF membrane targeted the removal of monomers, resulting in high RC for oligomers (0.95 for lactose/allolactose and 0.99 for GOS) and an expected increase in RC for glucose and galactose (0.56–0.58) due to reduction in pore size and membrane fouling, leaving about 5% of these monosaccharides in the final product. Despite this minor retention, the outcome is satisfactory for developing a commercial GOS ingredient.

### 3.2 ALPINA GOS have a prebiotic effect by inducing the production of higher levels of lactate and acetate and lowering the pH in MicroColon fermentation

After 20 hours of fermentation in the MicroColon model using two doses of ALPINA GOS, a clear pH lowering effect can be observed by both treatments (Figure 1A). The pH of the culture medium was lower in the treatment with ALPINA GOS compared to the non-treated (water) control (dashed line) and the control of MonoDiMix, which represents the mono- and disaccharide content present in the ALPINA GOS. Although the MonoDiMix also had a lowering effect on the pH, it was less pronounced than that of ALPINA GOS, indicating an additive effect of GOS beyond the contribution of the mono- and disaccharide mixture. Similarly, the scFOS control demonstrated a comparable pH-lowering effect to ALPINA GOS.

**Figure 1.**
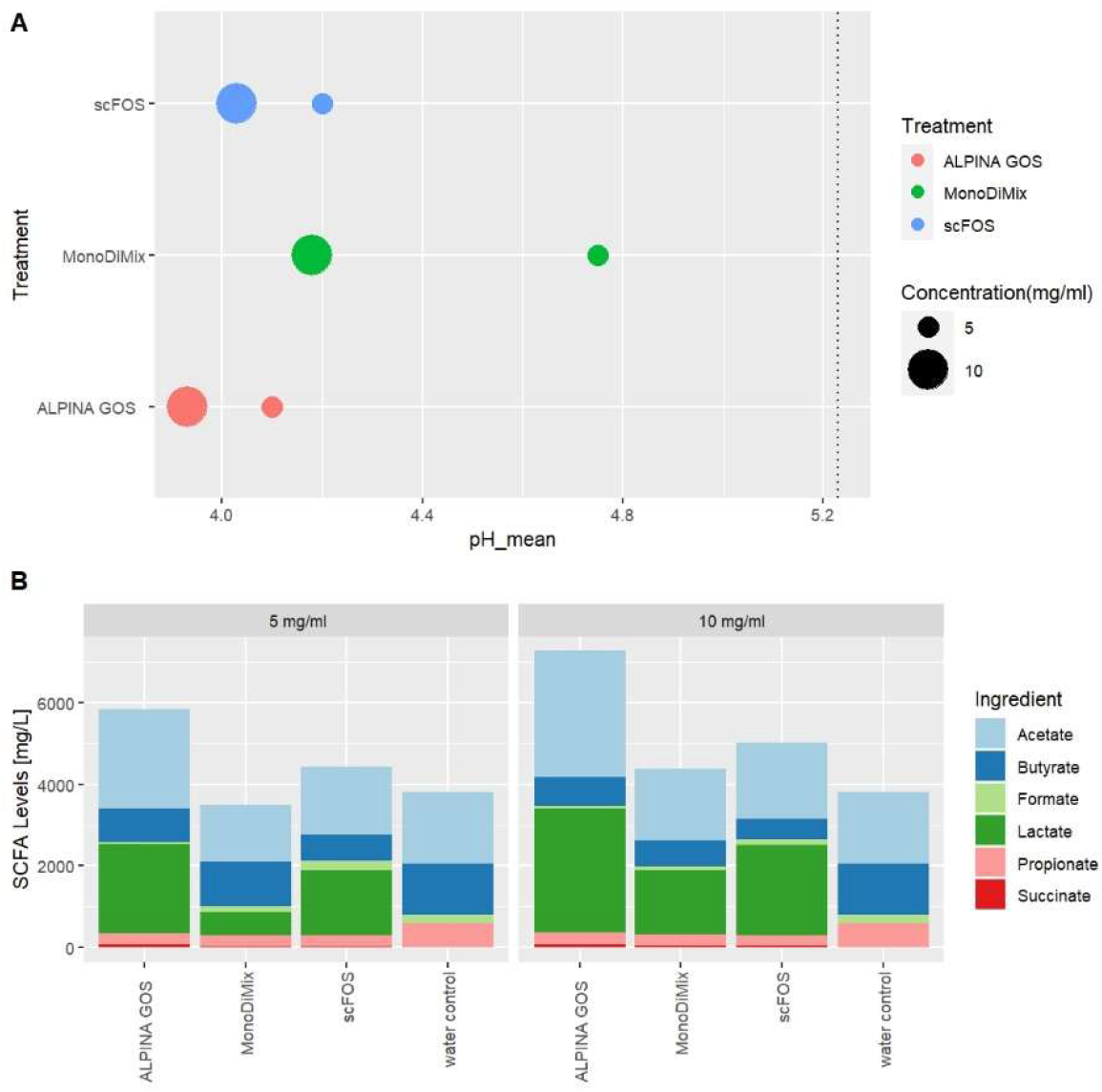
Final pH and Short Chain Fatty Acid composition after 20h of fermentation using 5 and 10 mg/ml concentration of ALPINA GOS, in the MicroColon experiment. **A.** pH of the water control is indicated by the dashed line. The circles representing each treatment are the mean of the pH values in the MicroColon simulations for 4 donors; the variability between donors is shown in SM 1A **B**. Each bar represents the mean values of SCFA abundance for the MicroColon simulations in each one of the treatments, for the 4 donors. Acetate and the sum of SCFA were significantly higher for the 5mg/mL dose of ALPINA GOS compared to the MonoDiMix (both p=0.0155; Friedman with Dunn’s posthoc test); the variability between donors can be found in SM 1B.

Notably, the 10 mg/mL dose of ALPINA GOS had a stronger and more pronounced effect than the 5 mg/ml dose (Figure 1; Supplementary Material SM1). The increased acidity in the ALPINA GOS treatments resulted from the fermentation of these oligosaccharides by the fecal microbiota, producing short-chain fatty acids (SCFAs) such as lactic and acetic acid (Figure 1B). The resulting lower pH creates a favorable environment for the growth of beneficial bacteria such us *Lactobacillus* and *Bifidobacterium*, while inhibiting the proliferation of pathogenic microorganisms (Macfarlane & Macfarlane, 2011; Roberfroid, 2007). Furthermore, an acidic environment enhances nutrient absorption and supports metabolic functions within the gut, contributing to overall intestinal health. These findings highlight the potential of ALPINA GOS as a prebiotic agent capable of modulating gut microbiota composition and function (Cani & Knauf, 2016; Morrison & Preston, 2016; Slavin, 2013).

Regarding the SCFA composition (Figure 1B) at 20h, treatment with 5 or 10 mg/ml of ALPINA GOS in the MicroColon model induced an increase in lactate and acetate levels. This effect was also observed, although to a lesser extent, with the MonoDiMix controls, compared to the water control. Similarly, control scFOS also induced an increase in acetate and lactate compared to the water control, as expected. Statistical analysis comparing the ALPINA GOS and MonoDiMix conditions using Friedman tests with Dunn’s posthoc tests, showed that acetate and the sum of SCFA were significantly higher for the 5mg/mL dose of ALPINA GOS (both p=0.0155). In contrast, butyrate production was (non-significantly) reduced in the GOS treatments, which aligns with the expectation that GOS primarily stimulate growth of non-butyrate-producing bacteria (Liu et al., 2017). Consistent with the pH results, the 10 mg/mL dose exhibited a stronger and more pronounced effect than the 5 mg/ml dose in overall SCFA production (Supplementary Material S1). Propionate, succinate and formate were not significantly stimulated compared to controls. Particularly for acetate, evidence suggests its potential to positively modulate host energy and substrate metabolism within the gut. Acetate has been described to elicit the secretion of gut hormones such as glucagon-like peptide-1 and peptide YY (Facchin et al., 2024; Hernández et al., 2019), while also promoting the increase in the relative abundance of beneficial butyrate producers such as *Faecalibacterium prausnitzii* and *Roseburia intestinalis*/*Eubacterium rectale* (Duncan et al., 2002).

These results demonstrated the prebiotic effect of ALPINA GOS, as evidenced by the increased production of lactate and acetate and a more pronounced reduction in pH compared to the MonoDiMix control. ALPINA GOS enhanced the production of SCFA, contributing to a more acidic gut environment, which supports gut health and promotes the growth of beneficial gut bacteria. These findings align with previous studies by Scott et al. (2013) (Scott et al., 2013) and Arnold et al. (2021) (Arnold et al., 2021), which demonstrated that dietary GOS modulates gut homeostasis, even during aging, by promoting microbiome changes that reduce intestinal permeability and increase mucus production. Moreover, it is well documented that the fermentation of GOS leads to the production of SCFAs and can promote the proliferation of beneficial bacteria beyond *Bifidobacteria*. For instance, Zou *et al* 2020 (Zou et al., 2020) reported a significant increase in the abundance of *Akkermansia_muciniphila* and *Ruminococcaceae* UCG 010, both effective SCFA producers, following GOS administration. Notably, a recent review by Yoo *et al* 2024 (Yoo et al., 2024) highlighted the vital role of GOS in modulating gut health, reducing inflammation, and enhancing immune responses, thereby offering broad health benefits.

### 3.3 The effects of ALPINA GOS on the adult gut microbiota fermentation processes denote a stimulation of beneficial bacteria, mostly *Bifidobacterium*

The prebiotic effect of GOS, as demonstrated by *in vitro* SCFA production (Figure 1B), was further corroborated through the compositional analysis of adult fecal microbiota following fermentation in the MicroColon model. After 20 hours of fermentation with ALPINA GOS by the gut microbiota of healthy adults, and compared to several controls (water, MonoDiMix, and a scFOS mixture), a clear bifidogenic effect was observed (Figure 2; Supplementary Material SM2). Specifically, Figures 2A and 2B show that the relative abundance of *Bifidobacterium* was (non-significantly) higher in the ALPINA GOS treatment compared to the MonoDiMix control (p=0.66). In addition to these changes, a strong individual signature of each donor was identified over time, as confirmed by PERMANOVA analysis at t20 (p=0.001; Figure 2C and SM3). This finding underscores the variability in microbiome composition among individuals and highlights the potential for personalized nutrition strategies to optimize health outcomes. By tailoring GOS supplementation to individual microbiome profiles, it may be possible to enhance gut health more effectively, as GOS selectively promotes beneficial bacteria that can vary significantly from person to person. These results align with emerging research that emphasizes the importance of personalized nutrition in improving health outcomes (Sarfraz et al., 2022).

**Figure 2.**
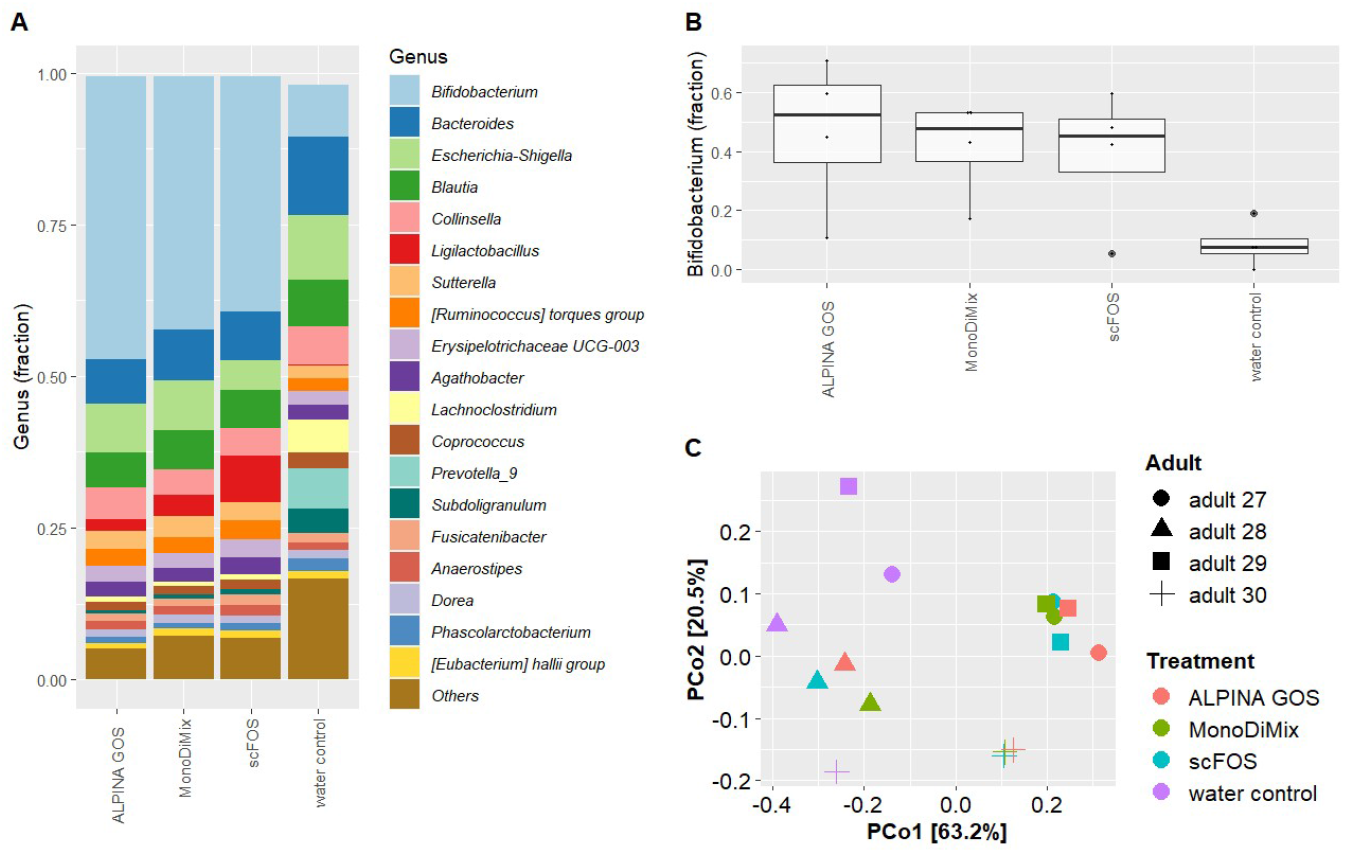
Gut microbiome profiling after 20h of fermentation with ALPINA GOS in the MicroColon model. **A.** Mean composition of the top-20 most abundant genera per treatment, (average of the 4 donors; the variability between donors is shown in SM 2A). **B**. Relative abundance of *Bifidobacterium* in each one of the treatments, for the 4 donors. **C**. Principal Coordinates analysis (weighted UniFrac) of the microbiome composition of the 4 donors.

The bifidogenic effect of GOS is well documented (Arnold et al., 2021; Liu et al., 2017; Mei et al., 2022; Mysore Saiprasad et al., 2023). Consistent with these findings, our results demonstrate that ALPINA GOS, a carbohydrate complex derived from whey permeate, stimulates the growth of lactate- and acetate-producing *Bifidobacterium* in the adult microbiota. The production of organic acids, such as lactate and acetate, not only lowers intestinal pH, creating an unfavorable environment for pathogens, but also serves as a critical energy source and as signaling molecules that promote cross-feeding interactions within the healthy microbiome. These cross-feeding mechanisms can foster the growth of other beneficial bacteria, such as *Roseburia*, which are known to contribute to eubiosis and enhance the overall health of the gut microbiota (Figure 2A; Supplementary Material SM2) (Pier et al., 2020; Portincasa et al., 2022).

Next, the addition of GOS in MicroColon simulations resulted in a decrease in microbial richness/evenness, but an increase in phylogenetic diversity in the adult microbiota (Figure 3). Shannon diversity, which accounts for both species richness and evenness, was lower in the ALPINA GOS, MonoDiMix, and scFOS conditions after 20 hours of fermentation compared to the water control. This reduction aligns with the dominance of *Bifidobacterium* observed in these conditions. Conversely, ALPINA GOS and scFOS slightly increased the phylogenetic diversity of the microbiota compared to the mono-/disaccharides and water controls. These results suggest that although evenness was reduced by ALPINA GOS and scFOS, phylogenetic diversity was maintained. High functional microbial diversity is generally considered beneficial for human health, as reduced diversity has been linked to diseases such as irritable bowel syndrome (IBS), inflammatory bowel disease (IBD), and obesity (Bäckhed et al., 2012).

**Figure 3.**
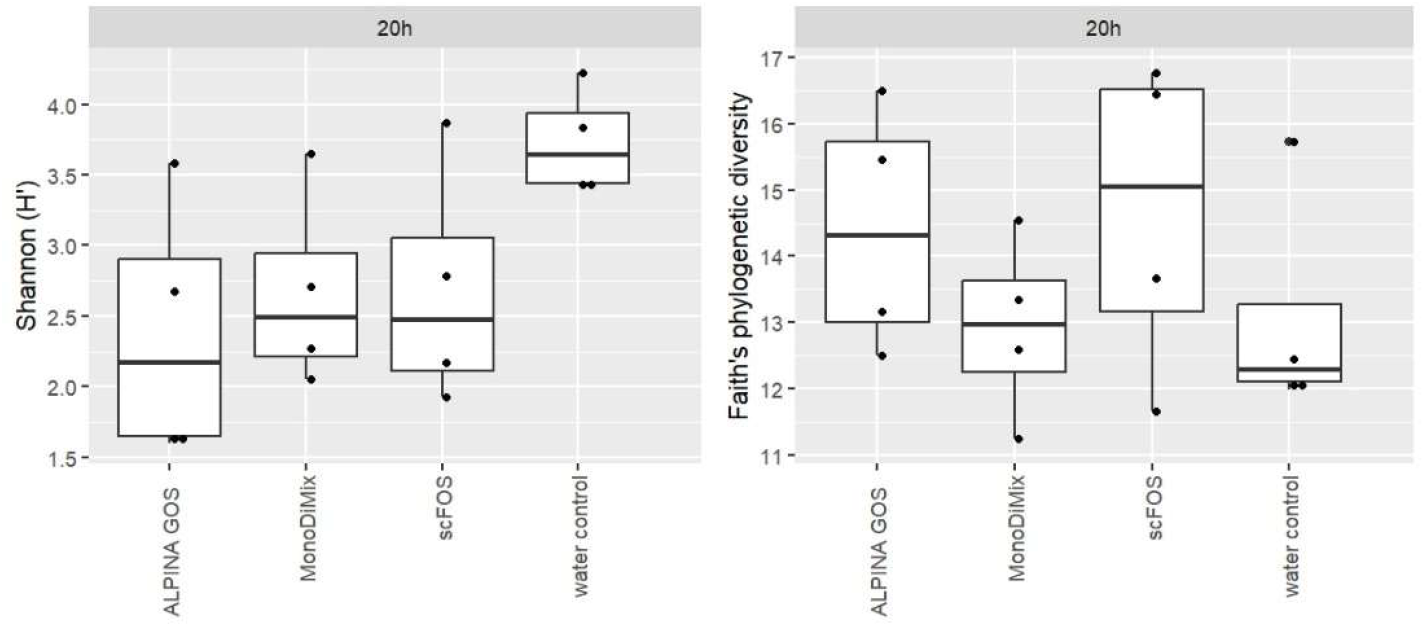
Alpha diversity, by Shannon index and Faith’s phylogenetic diversity after 20h of fermentation.

A comparative analysis between complex sugars such as GOS and simple sugars like those in the MonoDiMix treatment highlights the significant advantages of GOS. Complex sugars are recognized for their role in supporting healthy eating habits, while simple sugars are often avoided in nutritional recommendations due to their rapid metabolism, glucose-spiking effects, and limited availability for gut microbiota fermentation in the colon (Southgate, 1998). Additionally, there is evidence of negative effects associated with simple sugars reaching the colon in substantial amounts (Coker et al., 2021).

Current results, well supported by literature (Arnold et al., 2021; Liu et al., 2017), are particularly relevant as there is a critical need for high-quality evidence on the real effects of complex carbohydrates in enhancing specific beneficial microbial populations (Julien et al., 2023; Leeming et al., 2021). Robust evidence is essential to define the role of food-derived molecules, such as complex carbohydrates, in improving metabolic health through precise dietary recommendations. These effects are particularly notable when beneficial microorganisms contribute to eubiosis through diverse mechanisms, including the degradation of fructans, pectins, and other complex sugars (Pier et al., 2020; Portincasa et al., 2022).

## 4. Conclusions

This study clearly demonstrated that ALPINA GOS, produced through the enzymatic conversion of lactose in concentrated whey permeate, exerts beneficial effects on human gut microbial communities. Specifically, ALPINA GOS promotes an increase in the relative abundance of beneficial species, particularly *Bifidobacterium* spp., and elevates the production of lactic and acetic acids, and contributes to a lower pH in the gut environment. Although exploratory (n=4), this study presents robust data that clearly supports the bifidogenic effect of a GOS-rich ingredient obtained from an industrial by-product such as whey permeate. A wider study, *in vitro* and/or *in vivo*, would be necessary to validate these results and better understand the effect of this novel ingredient on the human gut microbiota and gut environment. Incorporating ALPINA GOS into new or existing formulations or products may enhance their potential gut health benefits.

## Acknowledgments

The authors thank Alpina Productos Alimenticios S.A.S. BIC for the financial support of this research. We also thank the fecal donors for their contribution to the study, and all the NIZO laboratory personnel who supported the execution of the study.

## Data availability statement

The data that supports the findings of this study are available in the supplementary material of this article.

## Ethics Statement

Human fecal samples used in this study were obtained from healthy adult volunteers who provided written informed consent prior to donation. All procedures were conducted in accordance with institutional ethical guidelines and the Declaration of Helsinki. No personally identifiable information was collected, and all samples were anonymized prior to analysis.

## References

Arnold, J. W., Roach, J., Fabela, S., Moorfield, E., Ding, S., Blue, E., Dagher, S., Magness, S., Tamayo, R., Bruno-barcena, J. M., & Azcarate-peril, M. A. (2021). The pleiotropic effects of prebiotic galacto-oligosaccharides on the aging gut. Microbiome, 9(31), 1–19. 10.1186/s40168-020-00980-0

Bäckhed, F., Fraser, C. M., Ringel, Y., Sanders, M. E., Sartor, R. B., Sherman, P. M., Versalovic, J., Young, V., & Finlay, B. B. (2012). Defining a healthy human gut microbiome: Current concepts, future directions, and clinical applications. Cell Host and Microbe, 12(5), 611–622. 10.1016/j.chom.2012.10.012

Bylund, G. (1995). Dairy processing handbook (T. AB (ed.); 1st ed.). Tetra Pak Processing Sustems AB.

Callahan, B. J., McMurdie, P. J., Rosen, M. J., Han, A. W., Johnson, A. J. A., & Holmes, S. P. (2016). DADA2: High-resolution sample inference from Illumina amplicon data. Nature Methods 2016 13:7, 13(7), 581–583. 10.1038/nmeth.3869

Canani, R. B., Costanzo, M. Di, Leone, L., Pedata, M., Meli, R., & Calignano, A. (2011). Potential beneficial effects of butyrate in intestinal and extraintestinal diseases. World Journal of Gastroenterology, 17(12), 1519–1528. 10.3748/wjg.v17.i12.1519

Cani, P. D., & Knauf, C. (2016). How gut microbes talk to organs: The role of endocrine and nervous routes. Molecular Metabolism, 5(9), 743–752. 10.1016/j.molmet.2016.05.011

Chen, G. Q., Qu, Y., Gras, S. L., & Kentish, S. E. (2023). Separation Technologies for Whey Protein Fractionation. Food Engineering Reviews, 15(3), 438–465. 10.1007/s12393-022-09330-2

Coker, J. K., Moyne, O., Rodionov, D. A., & Zengler, K. (2021). Carbohydrates great and small, from dietary fiber to sialic acids: How glycans influence the gut microbiome and affect human health. Gut Microbes, 13(1), 1–18. 10.1080/19490976.2020.1869502

Córdova, A., Astudillo, C., Giorno, L., Guerrero, C., Conidi, C., Illanes, A., & Cassano, A. (2016). Nanofiltration potential for the purification of highly concentrated enzymatically produced oligosaccharides. Food and Bioproducts Processing, 98, 50–61. 10.1016/j.fbp.2015.11.005

Coulier, L., Timmermans, J., Bas, R., Van den Dool, R., Haaksman, I., Klarenbeek, B., Slaghek, T., & Van Dongen, W. (2009). In-Depth Characterization of Prebiotic Galacto-oligosaccharides by a Combination of Analytical Techniques. Journal of Agricultural and Food Chemistry, 57, 8488–8495. 10.1021/jf902549e

Dixon, P. (2003). VEGAN, a package of R functions for community ecology. Journal of Vegetation Science, 14(6), 927–930. 10.1111/J.1654-1103.2003.TB02228.X

Duncan, S. H., Hold, G. L., Harmsen, H. J. M., Stewart, C. S., & Flint, H. J. (2002). Growth requirements and fermentation products of Fusobacterium prausnitzii, and a proposal to reclassify it as Faecalibacterium prausnitzii gen. nov., comb. nov. International Journal of Systematic and Evolutionary Microbiology, 52(6), 2141–2146. 10.1099/ijs.0.02241-0

Facchin, S., Bertin, L., Bonazzi, E., Lorenzon, G., De Barba, C., Barberio, B., Zingone, F., Maniero, D., Scarpa, M., Ruffolo, C., Angriman, I., & Savarino, E. V. (2024). Short-Chain Fatty Acids and Human Health: From Metabolic Pathways to Current Therapeutic Implications. Life, 14(5), 1–44. 10.3390/life14050559

Feng, Y. M., Chang, X. L., Wang, W. H., & Ma, R. Y. (2009). Separation of galacto-oligosaccharides mixture by nanofiltration. Journal of the Taiwan Institute of Chemical Engineers, 40(3), 326–332. 10.1016/j.jtice.2008.12.003

Fischer, C., & Kleinschmidt, T. (2018). Synthesis of Galactooligosaccharides in Milk and Whey: A Review. Comprehensive Reviews in Food Science and Food Safety, 17(3), 678–697. 10.1111/1541-4337.12344

Gommers, L. M. M., Ederveen, T. H. A., Wijst, J. Van Der, Overmars-Bos, C., Kortman, G. A. M., Boekhorst, J., Bindels, R., de Baaij, J., & Hoenderop, J. (2019). Low gut microbiota diversity and dietary magnesium intake are associated with the development of PPI-induced hypomagnesemia. The FASEB Journal, 33, 11235–11246. 10.1096/fj.201900839R

Griffin, I. J., Davila, P. M., & Abrams, S. A. (2002). Non-digestible oligosaccharides and calcium absorption in girls with adequate calcium intakes. British Journal of Nutrition, 87(S2), S187–S191. 10.1079/bjn/2002536

Grimaldi, R., Swann, J. R., Vulevic, J., Gibson, G. R., & Costabile, A. (2016). Fermentation properties and potential prebiotic activity of Bimuno® galacto-oligosaccharide (65 % galacto-oligosaccharide content) on in vitro gut microbiota parameters. British Journal of Nutrition, 116(3), 480–486. 10.1017/S0007114516002269

Hernández, M. A. G., Canfora, E. E., Jocken, J. W. E., & Blaak, E. E. (2019). The short-chain fatty acid acetate in body weight control and insulin sensitivity. Nutrients, 11(8). 10.3390/nu11081943

Hong, K. B., Jeong, M., Han, K. S., Hwan Kim, J., Park, Y., & Suh, H. J. (2015). Photoprotective effects of galacto-oligosaccharide and/or Bifidobacterium longum supplementation against skin damage induced by ultraviolet irradiation in hairless mice. International Journal of Food Sciences and Nutrition, 66(8), 923–930. 10.3109/09637486.2015.1088823

Julien, B., Julien, I. B., Paradis, S. C., Rochefort, G., Perron, J., Lamarche, B., Flamand, N., Marzo, V. Di, Veilleux, A., & Raymond, F. (2023). The diet rapidly and differentially affects the gut microbiota and host lipid mediators in a healthy population. Microbiome, 11(26), 1–16. 10.1186/s40168-023-01469-2

Kassambara, A. (2019). rstatix: Pipe-friendly framework for basic statistical tests. CRAN:Contributed Packages.

Katoh, K., Asimenos, G., & Toh, H. (2009). Multiple alignment of DNA sequences with MAFFT. Methods in Molecular Biology (Clifton, N.J.), 537, 39–64. 10.1007/978-1-59745-251-9_3

Kaur, H., Kaur, G., & Ali, S. A. (2024). Postbiotics Implication in the Microbiota-Host Intestinal Epithelial Cells Mutualism. Probiotics and Antimicrobial Proteins, 16, 443–458. 10.1007/s12602-023-10062-w

Kelly, C. J., Zheng, L., Campbell, E. L., Saeedi, B., Scholz, C. C., Bayless, A. J., Wilson, K. E., Glover, L. E., Kominsky, D. J., Magnuson, A., Weir, T. L., Ehrentraut, S. F., Pickel, C., Kuhn, K. A., Lanis, J. M., Nguyen, V., Taylor, C. T., & Colgan, S. P. (2015). Crosstalk between Microbiota-Derived Short-Chain Fatty Acids and Intestinal Epithelial HIF Augments Tissue Barrier Function. Cell Host & Microbe, 17(5), 662–671. 10.1016/J.CHOM.2015.03.005

Kembel, S. W., Cowan, P. D., Helmus, M. R., Cornwell, W. K., Morlon, H., Ackerly, D. D., Blomberg, S. P., & Webb, C. O. (2010). Picante: R tools for integrating phylogenies and ecology. Bioinformatics, 26(11), 1463–1464. 10.1093/BIOINFORMATICS/BTQ166

Kim, B., Kim, J. N., & Cerniglia, C. E. (2011). In Vitro Culture Conditions for Maintaining a Complex Population of Human Gastrointestinal Tract Microbiota. Journal of Biomedicine and Biotechnology, 838040, 10. 10.1155/2011/838040

Kortman, G. A. M., van Alen, I., Beerthuyzen, M., Lucas-van de Bos, E., van Schalkwijk, S., Staring, G., Jacobs, S., Hester, E. R., Scheithauer, T. P. M., & Hartog, A. (2023). Polyphenol-rich extracts from olive leaves modulate gut microbiota composition and metabolism in the in vitro MicroColon model, and show protective effects on the intestinal barrier function. 10th Beneficial Microbes Conference.

Krishnamurthy, H. K., Pereira, M., Bosco, J., George, J., Jayaraman, V., Krishna, K., Wang, T., Bei, K., & Rajasekaran, J. J. (2023). Gut commensals and their metabolites in health and disease. Frontiers in Microbiology, 14. 10.3389/fmicb.2023.1244293

Leeming, E. R., Louca, P., Gibson, R., Menni, C., Spector, T. D., & Le Roy, C. I. (2021). The complexities of the diet-microbiome relationship: advances and perspectives. Genome Medicine, 13, 1–14. 10.1186/s13073-020-00813-7

Litvak, Y., Byndloss, M. X., & Bäumler, A. J. (2018). Colonocyte metabolism shapes the gut microbiota. Science, 362(6418). 10.1126/SCIENCE.AAT9076/ASSET/1EF05AC3-94E6-4796-B35E-DD29B79C0FFD/ASSETS/GRAPHIC/362_AAT9076_FA.JPEG

Liu, F., Li, P., Chen, M., Luo, Y., Mujagond, P., Zheng, H., He, Y., Qi, Q., Long, H., Zhang, Y., Sheng, H., & Zhou, H. (2017). Fructooligosaccharide (FOS) and Galactooligosaccharide (GOS) Increase Bifidobacterium but Reduce Butyrate Producing Bacteria with Adverse Glycemic Metabolism in healthy young population. Scientific Reports, 7. 10.1038/s41598-017-10722-2

Liu, Y., Wang, J., & Wu, C. (2022). Modulation of Gut Microbiota and Immune System by Probiotics, Pre-biotics, and Post-biotics. Frontiers in Nutrition, 8(January), 1–14. 10.3389/fnut.2021.634897

Lozupone, C., & Knight, R. (2005). UniFrac: a new phylogenetic method for comparing microbial communities. Applied and Environmental Microbiology, 71(12), 8228–8235. 10.1128/AEM.71.12.8228-8235.2005

Macfarlane, G.T., Macfarlane, S., & Gibson, G. R. (1998). Validation of a Three-Stage Compound Continuous Culture System for Investigating the Effect of Retention Time on the Ecology and Metabolism of Bacteria in the Human Colon. Microbial Ecology, 35, 180–187. 10.1007/s002489900072

Macfarlane, George T, & Macfarlane, S. (2011). Fermentation in the Human Large Intestine Its Physiologic Consequences and the Potential Contribution of Prebiotics. Journal of Clinical Gastroenterology, 45, S120–S127. 10.1097/mcg.0b013e31822fecfe

Mei, Z., Yuan, J., & Li, D. (2022). Biological activity of galacto-oligosaccharides: A review. Frontiers in Microbiology, 13(September), 1–7. 10.3389/fmicb.2022.993052

Michelon, M., Manera, A. P., Carvalho, A. L., & Maugeri Filho, F. (2014). Concentration and purification of galacto-oligosaccharides using nanofiltration membranes. International Journal of Food Science and Technology, 49(8), 1953–1961. 10.1111/ijfs.12582

Morrison, D. J., & Preston, T. (2016). Formation of short chain fatty acids by the gut microbiota and their impact on human metabolism. Gut Microbes, 7(3), 189–200. 10.1080/19490976.2015.1134082

Mysore Saiprasad, S., Moreno, O. G., & Savaiano, D. A. (2023). A Narrative Review of Human Clinical Trials to Improve Lactose Digestion and Tolerance by Feeding Bifidobacteria or Galacto-Oligosacharides. Nutrients, 15(16). 10.3390/nu15163559

Nobre, C., Santos, M. J., Dominguez, A., Torres, D., Rocha, O., Peres, A. M., Rocha, I., Ferreira, E. C., Teixeira, J. A., & Rodrigues, L. R. (2009). Comparison of adsorption equilibrium of fructose, glucose and sucrose on potassium gel-type and macroporous sodium ion-exchange resins. Analytica Chimica Acta, 654(1), 71–76. 10.1016/j.aca.2009.06.043

Olivares-Tenorio, M.-L., Orrego, D., Klotz-Ceberio, B.-F., Palanca, C., & Tortajada-serra, M. (2022). Galactooligosaccharides: Food technological applications, prebiotic health benefits, microbiome modulation, and processing considerations. JSFA Reports, 4(7), 1–13. 10.1002/jsf2.92

Orrego, D., & Klotz-Ceberio, B. (2022). Enzymatic Synthesis of Galacto-Oligosaccharides from Concentrated Sweet Whey Permeate and Its Application in a Dairy Product. Applied Sciences 2022, Vol. 12, Page 10229, 12(20), 10229. 10.3390/APP122010229

Pathan, S., Glover, M., Ryan, J., & Shih, D. Q. (2024). The Role of Inulin in Human Health and Sustainable Food Applications. IntechOpen. 10.5772/intechopen.1007006

Pier, M., Guarino, L., Altomare, A., Emerenziani, S., Di Rosa, C., Ribolsi, M., Balestrieri, P., Iovino, P., Rocchi, G., & Cicala, M. (2020). Mechanisms of Action of Prebiotics and Their Effects on Gastro-Intestinal Disorders in Adults. Nutrients, 12(4), 1–24. 10.3390/nu12041037

Portincasa, P., Bonfrate, L., Vacca, M., Angelis, M. De, Farella, I., Lanza, E., Khalil, M., Wang, D. Q., Sperandio, M., & Di Ciaula, A. (2022). Gut Microbiota and Short Chain Fatty Acids: Implications in Glucose Homeostasis. International Journal of Molecular Science, 23(3). 10.3390/ijms23031105

Price, M. N., Dehal, P. S., & Arkin, A. P. (2010). FastTree 2 – Approximately Maximum-Likelihood Trees for Large Alignments. PLOS ONE, 5(3), e9490. 10.1371/JOURNAL.PONE.0009490

Quast, C., Pruesse, E., Yilmaz, P., Gerken, J., Schweer, T., Yarza, P., Peplies, J., & Glöckner, F. O. (2013). The SILVA ribosomal RNA gene database project: improved data processing and web-based tools. Nucleic Acids Research, 41(Database issue). 10.1093/NAR/GKS1219

R Core, T. (2023). R: A Language and Environment for Statistical Computing. R Foundation for Statistical Computing. https://www.r-project.org/

Roager, H. M., & Licht, T. R. (2018). Microbial tryptophan catabolites in health and disease. Nature Communications, 9(1), 1–10. 10.1038/s41467-018-05470-4

Roberfroid, M. (2007). Prebiotics: The Concept Revisited. The Journal of Nutrition, 137, 830S–837S. 10.1093/jn/137.3.830S

Roy, D., Ye, A., Moughan, P. J., & Singh, H. (2020). Composition, Structure, and Digestive Dynamics of Milk From Different Species—A Review. Frontiers in Nutrition, 7(October), 1–17. 10.3389/fnut.2020.577759

Sangwan, V., Tomar, S. K., Singh, R. R. B., Singh, A. K., & Ali, B. (2011). Galactooligosaccharides: Novel Components of Designer Foods. Journal of Food Science, 76(4). 10.1111/j.1750-3841.2011.02131.x

Sarfraz, M. H., Shahid, A., Asghar, S., Aslam, B., Ashfaq, U. A., Raza, H., Prieto, M. A., Simal-Gandara, J., Barba, F. J., Rajoka, M. S. R., Khurshid, M., & Nashwan, A. J. (2022). Personalized nutrition, microbiota, and metabolism: A triad for eudaimonia. Frontiers in Molecular Biosciences, 9(October), 1–15. 10.3389/fmolb.2022.1038830

Schmidt, C. M., Mailänder, L. K., & Hinrichs, J. (2019). Fractionation of mono- and disaccharides via nanofiltration: Influence of pressure, temperature and concentration. Separation and Purification Technology, 211(October 2018), 571–577. 10.1016/j.seppur.2018.10.024

Scott, K. P., Martin, J. C., Duncan, S. H., & Flint, H. J. (2013). Prebiotic stimulation of human colonic butyrate-producing bacteria and bifidobacteria, in vitro. FEMS Microbiology Ecology, 87(1), 30–40. 10.1111/1574-6941.12186

Senchukova, M. A. (2023). Microbiota of the gastrointestinal tract: Friend or foe? World Journal of Gastroenterology, 29(1), 19–42. 10.3748/wjg.v29.i1.19

Shkembi, B., & Huppertz, T. (2023). Glycemic Responses of Milk and Plant-Based Drinks: Food Matrix Effects. Foods, 12(3), 1–18. 10.3390/foods12030453

Slavin, J. (2013). Fiber and prebiotics: Mechanisms and health benefits. Nutrients, 5(4), 1417–1435. 10.3390/nu5041417

Southgate, D. A. T. (1998). How much and what classes of carbohydrate reach the colon. European Journal of Cancer Prevention, 7(2), S81–82. 10.1097/00008469-199805000-00014

Souza, A. F. C. e., Gabardo, S., & Coelho, R. de J. S. (2022). Galactooligosaccharides: Physiological benefits, production strategies, and industrial application. Journal of Biotechnology, 359(September), 116–129. 10.1016/j.jbiotec.2022.09.020

Tzortzis, G., & Vulevic, J. (2009). Galacto-Oligosaccharide Prebiotics. In D. Charalampopoulos & R. A. Rastall (Eds.), Prebiotics and Probiotics Science and Technology (pp. 207–244). Springer New York, NY. 10.1007/978-0-387-79058-9_7

Van Leusen, E., Torringa, E., Groenink, P., Kortleve, P., Geene, R., Schoterman, M., & Klarenbeek, B. (2014). Industrial Applications of Galactooligosaccharides. In F. J. Moreno & M. L. Sanz (Eds.), Food Oligosaccharides: Production, Analysis and Bioactivity (pp. 470–491). John Wiley & Sons, Ltd. 10.1002/9781118817360.ch25

Venegas, D. P., De La Fuente, M. K., Landskron, G., González, M. J., Quera, R., Dijkstra, G., Harmsen, H. J. M., Faber, K. N., & Hermoso, M. A. (2019). Short chain fatty acids (SCFAs)mediated gut epithelial and immune regulation and its relevance for inflammatory bowel diseases. Frontiers in Immunology, 10. 10.3389/fimmu.2019.00277

Wang, K., Duan, F., Sun, T., Zhang, Y., & Lu, L. (2023). Galactooligosaccharides: Synthesis, metabolism, bioactivities and food applications. Critical Reviews in Food Science and Nutrition, 1–17. 10.1080/10408398.2022.2164244

Wickham, H. (2011). ggplot2. Wiley Interdisciplinary Reviews: Computational Statistics, 3(2), 180–185. 10.1002/WICS.147

Yoo, S., Jung, S., Kwak, K., & Kim, J.-S. (2024). The Role of Prebiotics in Modulating Gut Microbiota: Implications for Human Health. International Journal of Molecular Sciences, 25(9). 10.3390/ijms25094834

Zhang, Z., Zhang, H., Chen, T., Shi, L., Wang, D., & Tang, D. (2022). Regulatory role of short-chain fatty acids in inflammatory bowel disease. Cell Communication and Signaling, 20(1), 1–10. 10.1186/s12964-022-00869-5

Zou, Y., Wang, J., Wang, Y., Peng, B., Liu, J., Zhang, B., Lv, H., & Wang, S. (2020). Protection of Galacto-Oligosaccharide against E. coli O157 Colonization through Enhancing Gut Barrier Function and Modulating Gut Microbiota. Foods, 9(11). 10.3390/foods9111710

