## Supplementary Material 3 for "Prebiotic ALPINA GOS produced from whey permeate has a bifidogenic effect on the adult fecal microbiota *in vitro*, including stimulation of organic acids production"

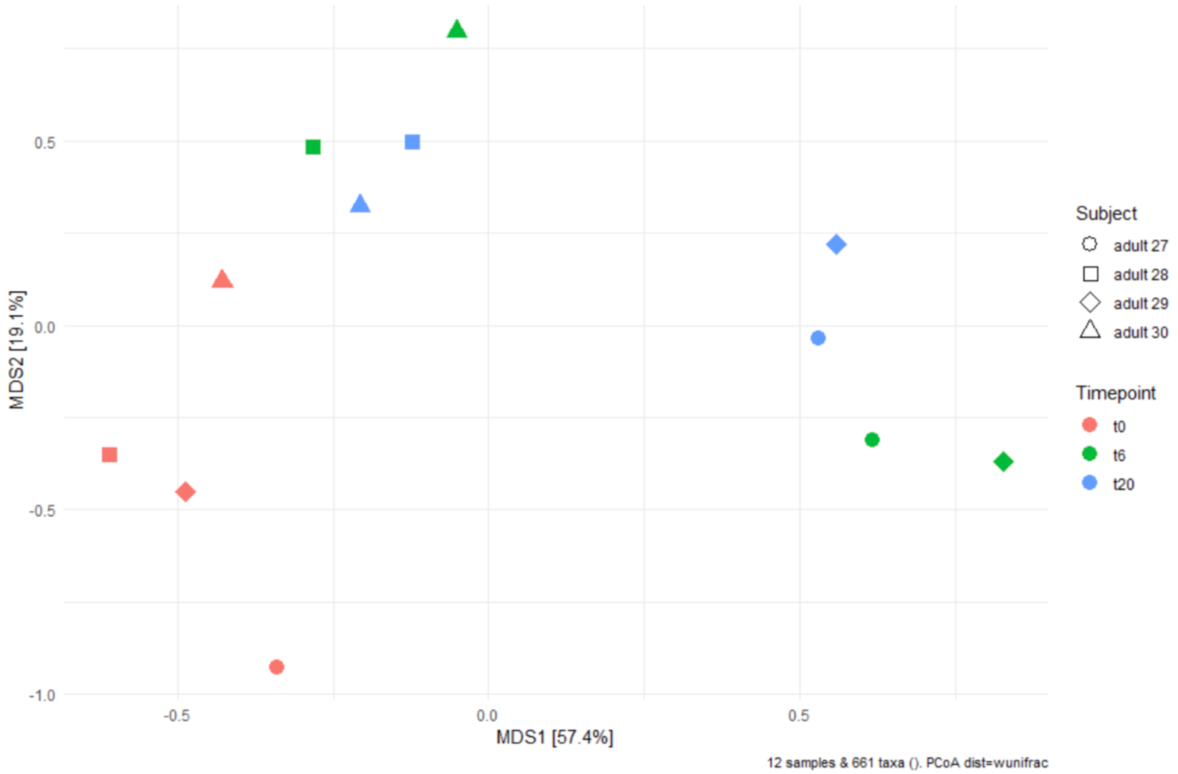


**SM3.** PCoA of weighted UniFrac distances between the untreated (water) control samples. In the MicroColon model, microbiota composition of the individual donors changed over time, but individual donor signatures remained.
